## Supplementary Appendix for "Microtubule re-organization during female meiosis in *C. elegans*"

### 1 Mathematical model for microtubule length distributions

The distribution $\psi(\ell)$ of microtubule lengths $\ell$ is set by the microtubule growth velocity $v_g$, their turnover rate $r$, and the rate at which microtubules are severed by proteins like Katanin $\kappa$. It obeys

$$\partial_t \psi = -\partial_\ell (v_g \psi) - (r + \kappa \ell)\psi + \kappa(1+\alpha) \int_\ell^\infty d\ell' \psi(\ell'), \tag{1}$$

Note that the dimensions of $\kappa$ is per time per length, and thus the longer microtubules are more likely to get cut. The dimensionless parameter $\alpha$, which can take values from zero to one, determines the likelihood that a new microtubule plus end created by severing is stable. After some basic algebra, we find that the steady states of Eq. (1) obey

$$v_g \partial_\ell^2 \psi + (r + \kappa\ell)\partial_\ell \psi + (2+\alpha)\kappa\psi = 0 \tag{2}$$

with the boundary conditions

$$v_g \psi(0) = rN - \kappa\alpha N \langle \ell \rangle , \tag{3}$$

$$\partial_\ell v_g \psi|_{\ell=0} = -r\psi(0) + \kappa(1+\alpha)N, \tag{4}$$

which state that the creation of new microtubules by nucleation and severing balances the loss of microtubules by turnover, such that the total number of microtubules $N$ stays a constant. Here $\langle \ell \rangle = \frac{1}{N}\int_0^\infty d\ell' \psi(\ell')\ell'$ denotes the average microtubule length.

For given parameters $r, \kappa, \alpha$ we can solve for the microtubule length $\psi(\ell)$ distribution numerically. Our custom written Code uses second order finite differences and is available from the authors upon reasonable request.

### 2 Bayesian Inference method for the relative importance of cutting and catastrophe

We seek to infer the probability distribution $P(model|data)$ of the model parameters $(r, \alpha, \kappa)$, from the electron tomography data. This can be done using Bayes' formula

$$P(model|data) = P(data|model)P(model)/P(data), \tag{5}$$

where $P(data|model)$ is the likelihood of the observed data given the model, $P(model)$ is the prior, and $P(data)$ in the finally marginalized likelihood. Since $P(data)$ is in general not easily calculated we use Markov Chain Monte Carlo sampling - specifically the toolbox (`https://github.com/dfn/emcee`) - to approach this problem. This has the advantage that we need not provide and expression for $P(data)$. We next discuss the expressions that we use for $P(data|model)$ and $P(model)$, respectively.

#### 2.1 Calculating $P(data|model)$

This term calculates the likelihood that the measured data (the observed microtubule lengths in our case) are a draw from the probability distribution function predicted by the model. To calculate this term we first solve Eq.2 with the model parameters and obtain the probability distribution $p(\ell) = \psi(\ell)/\int_0^\infty d\ell\psi(\ell)$, which characterizes the length distribution in the model.

To obtain the likelihood of the data given the model, we sort the data in to 256 bins of equal size distributed from length zero to 1.1 times the length of the longest microtubule seen in experiment. We then use $p(\ell)$ to determine the expectation value $\lambda_i$ for the number of microtubules found in bin $i$. (Note that we constrained the overall nucleation rate in the model to the value which would make the experimentally observed number of microtubules equal to the expected total number of microtubules.) With this the probability of finding $m_i$ microtubules in bin $i$ is Poisson distributed, and thus

$$P(data|model) = \sum(i)\frac{\lambda_i^{m_i}e^{-\lambda_i}}{m_i!}. \tag{6}$$

Note that during this procedure we ignore data for microtubules that are shorter than 150$n$m long, since we suspect that many of these are artefacts from the tomography reconstruction method, rather than real data points.

#### 2.2 Choosing a prior $P(model)$

We seek to not over-constrain our search space and thus use a so called uninformative prior. This means that we assign all models within the parameter ranges $\bar{r} > 0$, $\bar{\kappa} > 0$ and $0 < \alpha < 1$ the same a-priori likelihood.

#### 2.3 Test on generated test data

To test our inference scheme, we generated data sets from known distributions. We generated data sets that have comparable numbers and length of distributions as our experimental data and ran our inference scheme. In Figs . . . we show the results of this procedure, and compare against ground truth, for one case without and one case with cutting.

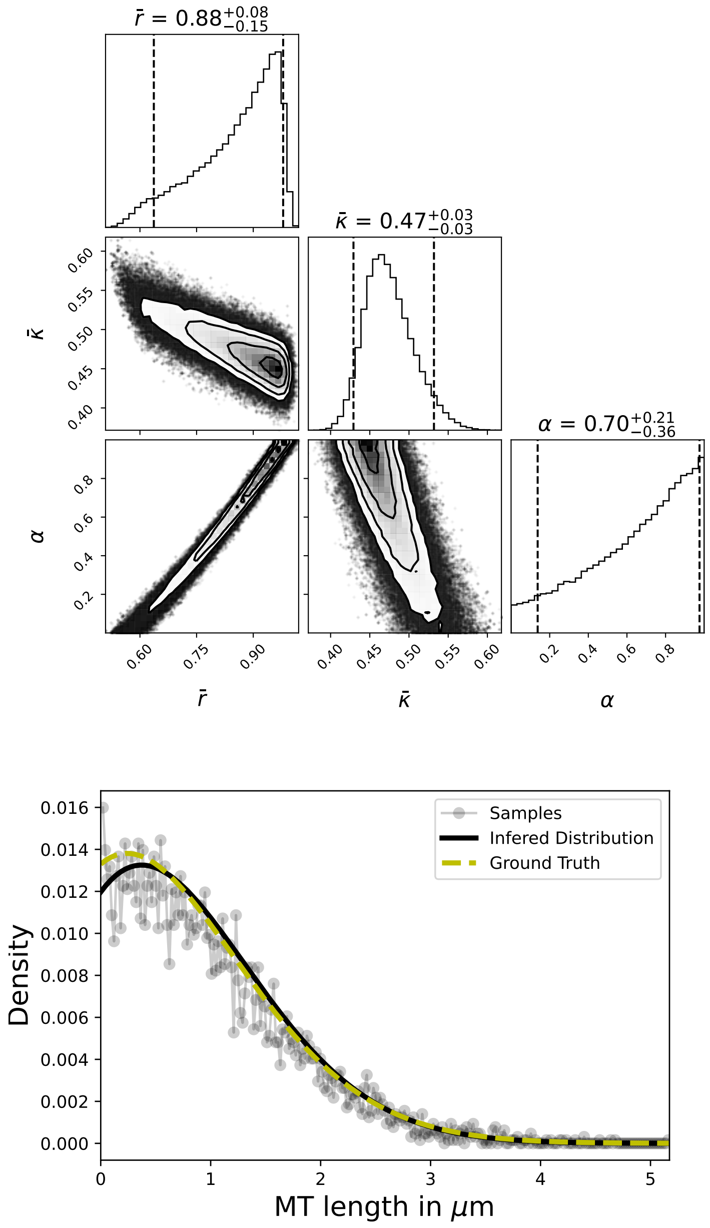


Figure 1: Results of inference vs ground thruths on artificial data, Case with cutting.

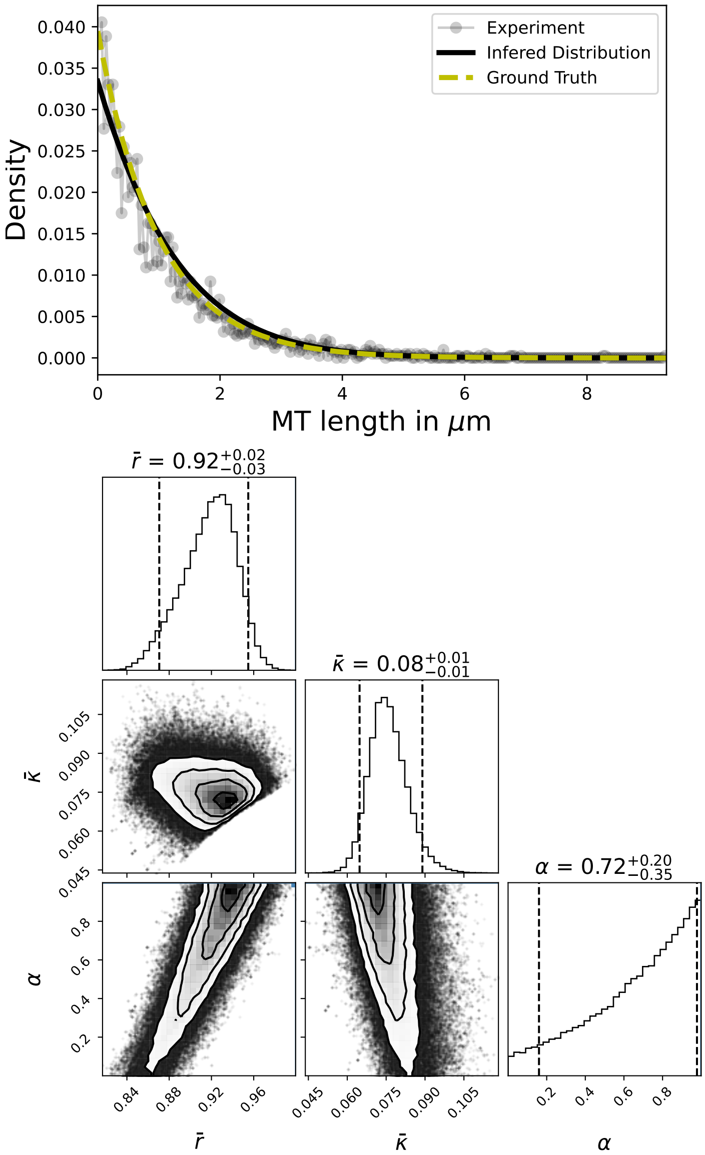


Figure 2: Results of inference vs ground thruths on artificial data, Case without cutting.
